# Delivery of recombinant SARS-CoV-2 envelope protein into human cells

**DOI:** 10.1101/2021.02.18.431684

**Authors:** James M. Hutchison, Ricardo Capone, Dustin D. Luu, Arina Hadziselimovic, Wade D. Van Horn, Charles R. Sanders

**Author notes:** These authors contributed equally (James M. Hutchison, Ricardo Capone).

## Abstract

SARS-CoV-2 envelope protein (S2-E) is a conserved membrane protein that is essential to coronavirus assembly and budding. Here, we describe the recombinant expression and purification of S2-E into amphipol-class amphipathic polymer solutions. The physical properties of amphipols underpin their ability to solubilize and stabilize membrane proteins without disrupting membranes. Amphipol delivery of S2-E to pre-formed planar bilayers results in spontaneous membrane integration and formation of viroporin ion channels. Amphipol delivery of the S2-E protein to human cells results in membrane integration followed by retrograde trafficking to a location adjacent to the endoplasmic reticulum-to-Golgi intermediate compartment (ERGIC) and the Golgi, which are the sites of coronavirus replication. Delivery of S2-E to cells enables both chemical biological approaches for future studies of SARS-CoV-2 pathogenesis and development of “Trojan Horse” anti-viral therapies. This work also establishes a paradigm for amphipol-mediated delivery of membrane proteins to cells.

## Introduction

The severe acute respiratory syndrome 2 virus (SARS-CoV-2) became a focal point of science and society in 2020. It is to be hoped that the ongoing vaccine development and delivery program will soon allow the world to return to an approximation of normalcy (1,2). However, previous coronavirus (CoV) epidemics, including Middle East respiratory syndrome (MERS) (3) and SARS (4) from 2002-2003 foretell that future CoV zoonotic events (5) are likely to afflict humankind. Fundamental studies of the molecular underpinnings of CoVs may help to mitigate the current and future pandemics.

Within CoVs, there are four critically conserved structural proteins (6,7), each of which is of possible therapeutic importance due to their essential functions (8), Among these is the SARS-CoV-2 envelope (E) protein. The E protein is a single-pass transmembrane protein whose roles in pathogenesis are incompletely understood (9). However, its importance is highlighted by cellular studies showing that the CoV E and M proteins alone are sufficient to produce a budding virus-like particle (VLP) (10–12). Moreover, deletion of E drastically lowers viral fitness (13–15) and growing evidence suggests that E is directly responsible for acute respiratory distress syndrome (ARDS) occurring in conjunction with CoV infections (16). E is highly expressed in infected cells, but only a small fraction is incorporated into mature viral particles, implying functions beyond its role as a mature capsid structural protein (17). Supporting this idea, the E protein is known to populate both monomer and oligomer forms *in vivo* (18). Most biophysical measurements have focused on the homopentamer form that functions as a cation-selective ion channel (19–22), which is analogous to a well-studied and validated drug target, the influenza M2 protein (23,24).

A distinct feature of coronavirus assembly is that their nascent particles bud into the lumen of the endoplasmic reticulum-to-Golgi intermediate compartments (ERGIC) in cells (25). The E protein is critical to viral maturation (10,17,26). Localization of SARS E to these membranes is remarkably stringent, likely a consequence of Golgi-targeting motifs present in the E protein (26). Since E functions in multiple roles that are critical to viral fitness (27–29), it is desirable to develop methods to further characterize key pathogenic mechanisms. Current methods to study the E protein in mammalian cells are reliant on transfection of genetic material encoding the protein into cells and its subsequent transcription and/or translation. Here, we sought to develop a robust method for exogenous delivery of purified SARS-CoV-2 envelope protein (S2-E) into cells to enable chemical biological methods for studies of S2-E function and to facilitate novel COVID therapies.

## Results and Discussion

We developed a straightforward bacterial expression and purification protocol that yields ~100 μg/L of 90-95% pure fulllength S2-E under conditions in which it is bereft of detergent and lipid, with its aqueous solubility being maintained by complexation with the zwitterionic amphipol PMAL-C8 (30,31) (**Fig. S1** and Supporting Material and Methods). This purification protocol has been streamlined to a single gravity column and does not require a FPLC or ultracentrifuge. Once purified into lipid/detergent-free amphipol solution, the S2-E/amphipol complexes remain stable and soluble in aqueous solution even following removal of excess uncomplexed amphipols. Amphipols are a class of amphipathic polymers that exhibit weak detergent properties, in that they can solubilize and stabilize the native membrane protein folds, but cannot solubilize or even permeabilize membranes (32,33). Additionally, some amphipols are well tolerated by animals (34) and have been used in Chlamydia vaccine development (35,36) because they do not elicit the production of anti-amphipol antibodies (37).

Planar lipid bilayer electrophysiology was used to test if amphipols could deliver the S2-E protein to a membrane environment to form ion channels without otherwise disrupting the lipid bilayer (**Fig. 1*A***). As expected, amphipol-based S2-E delivery resulted in ion channel activity that is consistent with previous SARS-CoV-1 E (38) and preliminary S2-E (39) channel measurements in terms of current amplitudes, sodium cation selectivity, and open probabilities. (**Figs. 1*B,C*, and S2**, and Supporting Materials and Methods). The S2-E-dependent currents and similarity to other planar bilayer measurements support that the idea that S2-E is released spontaneously from the amphipol into membranes. The bilayer integrity during amphipol delivery and exposure was monitored through membrane capacitance measurements. The bilayers remained stable throughout the recordings with an average value of 58 ± 3 pF. These results demonstrate that recombinant S2-E can be delivered into pre-formed lipid bilayers using amphipols, where the protein not only inserts into the bilayers, but also retains ion channel function, without significantly compromising the bilayer integrity.

**Figure 1.**
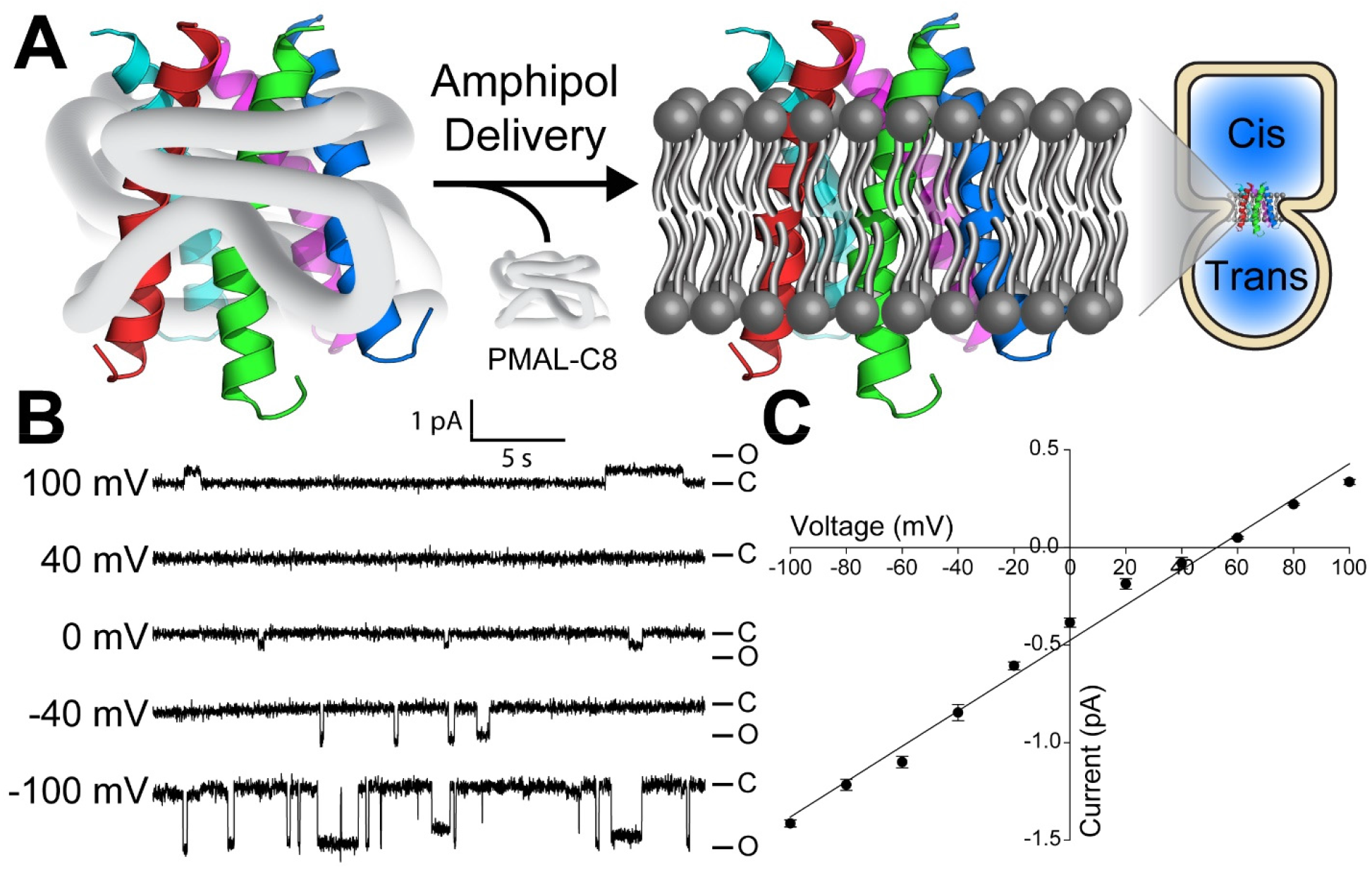
Functional delivery of SARS-CoV-2 envelope protein from amphipol complexes to planar lipid bilayers. **(A)** Schematic of SARS-CoV-2 envelope protein (S2-E) delivered using amphipols for membrane protein insertion into planar lipid bilayers. **(B)** Representative single-channel current recordings of PMAL-C8 amphipol-delivered S2-E as a function of transmembrane electrical potential show ion channel activity in POPC:POPE (3:1) planar bilayers, where S2-E fluctuates between closed (C) and open (O) states. **(C)** The S2-E current-voltage relationship identifies a conductance of 9.0 ± 0.3 pS and a reversal potential of 53 ± 3 mV in an asymmetric NaCl buffer, indicative of cation selectivity. Data represent three replicates. Error bars are SEM from the three distinct amphipol delivery experiments on different days.

We next tested whether S2-E can be delivered from amphipol complexes to the membranes of human cells. To this end, S2-E was irreversibly tagged with the fluorophore nitrobenzoxadiazole (NBD) to form S2-E-NBD. This allowed us to track the time course of delivery of S2-E into HeLa cells using confocal microscopy. As shown in **Figs. 2 and S5**, the S2-E-NBD protein was delivered from amphipol complexes to HeLa cell membranes, with all cells exhibiting NBD signal within 30 min (**Fig. 2*B***). **Fig. 2*C-F*** shows the 8 hour progression of the S2-E-NBD protein from the plasma membranes to a predominately perinuclear intracellular location. After 16-18 h nearly all the S2-E was observed in the vicinity of the nucleus, with a clear focal area on one side of the nuclear compartment rather than being evenly distributed, ring-like, around the entire nucleus (**Fig. 2*G,H***). Delivered S2-E was typically more diffuse at early time points but becomes punctate as it traffics to the perinuclear space.

**Figure 2.**
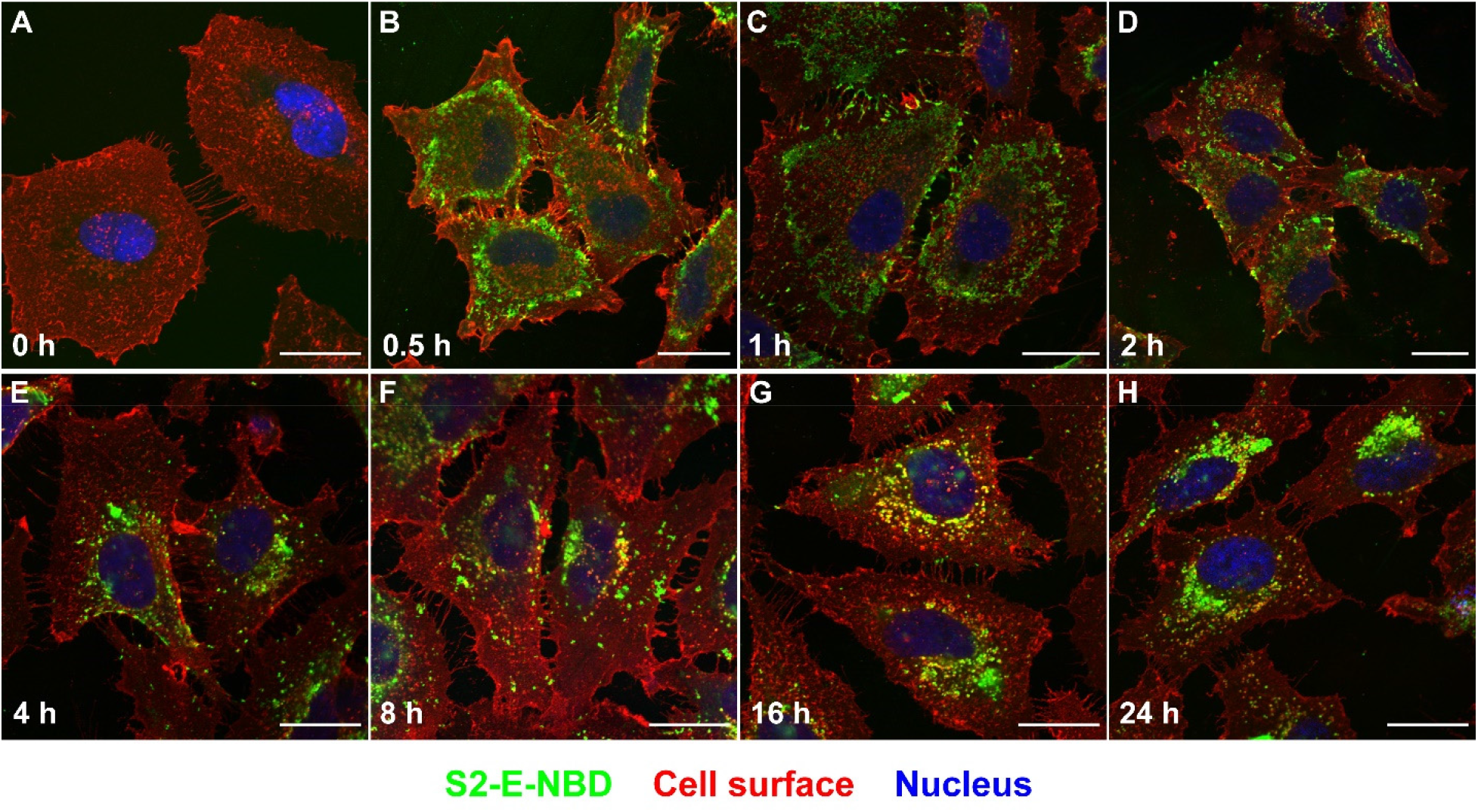
Uptake of amphipol delivered SARS CoV-2 E protein by cells and subsequent intracellular trafficking of the protein. Representative confocal microscopy images of HeLa cells at various time points following treatment with amphipol-complexed 2.5μM S2-E labeled with NBD (S2-E-NBD). Color markers are: green, S2-E-NBD; red, cell membrane (WGA-AF555); blue, cell nucleus (DRAQ5). **(A)** is the untreated (0 μM) sample and 0 h time point, **(B)** is cells 0.5 h after treatment, **(C)** is following 1 h, **(D)** 2 h, **(E)** 4 h, **(F)** 8 h, **(G)** 16 h, and **(H)** 24 h. The S2-E-NBD signal migrates from the cell plasma membrane (see panel B), towards the perinuclear space (see panels G and H). Time course experiments, using the same cell markers were independently repeated 3 times using 3 different S2-E-NBD preparations. All scale bars are 25 μm. See further details in the Supporting Information Materials and Methods and Fig. S5.

The amount of S2-E signal in cells was dependent on the applied amphipol/S2-E “dose” and no obvious cell toxicity was observed until a concentration of 10 μM S2-E in the culture was reached (**Figs. S3 and S4**). To ensure that we were microscopically tracking intact S2-E instead of dye freed from full length S2-E by degradation, we confirmed the S2-E localization following cell fixation and permeabilization with a polyclonal anti-S2-E antibody (**Fig. S3**). The same **Fig. S3** Western blot data also rules out the possibility that the tracked NBD fluorescence could arise from a minor impurity in our S2-E-NBD samples. While amphipols have previously been reported to deliver select membrane proteins to artificial lipid bilayers (30,40), this study represents the first use of amphipols to deliver a protein to live mammalian cells. Elucidation of the pathway(s) taken by the S2-E protein to dissociate from its soluble amphipol complex to then insert into the membrane to adopt a transbilayer configuration will require further study.

We also examined possible delivery of S2-E from amphipol solutions into SW1573 human alveolar cells, a COVID-19-relevant cell line (41). We observed (**Fig. S4**) that S2-E is indeed taken up by these cells and subject to the same cell surface-to-perinuclear “retrograde” trafficking as seen in HeLa cells.

During viral replication, most E protein is retained at the Golgi/ERGIC regions. S2-E retention is important to virion assembly because CoVs assemble and bud from the Golgi/ERGIC space before being secreted. The fact that S2-E retrograde traffics proximal to one side the nucleus (**Fig. 2*G,H***) is consistent with its localization at or near the Golgi/ERGIC compartments. To gain further insight into the final cellular location of S2-E we used organelle-specific monoclonal antibodies to pinpoint the locations of the Golgi and ERGIC relative to delivered S2-E. At later timepoints after initial delivery, S2-E was typically seen to concentrate in the area surrounding the Golgi, but not within the Golgi, (**Figs. 3*A-C* and S6**). In like manner, S2-E was seen to locate proximal to the cytosol-facing side of the ERGIC (see **Fig. 3*D-F***).

**Figure 3.**
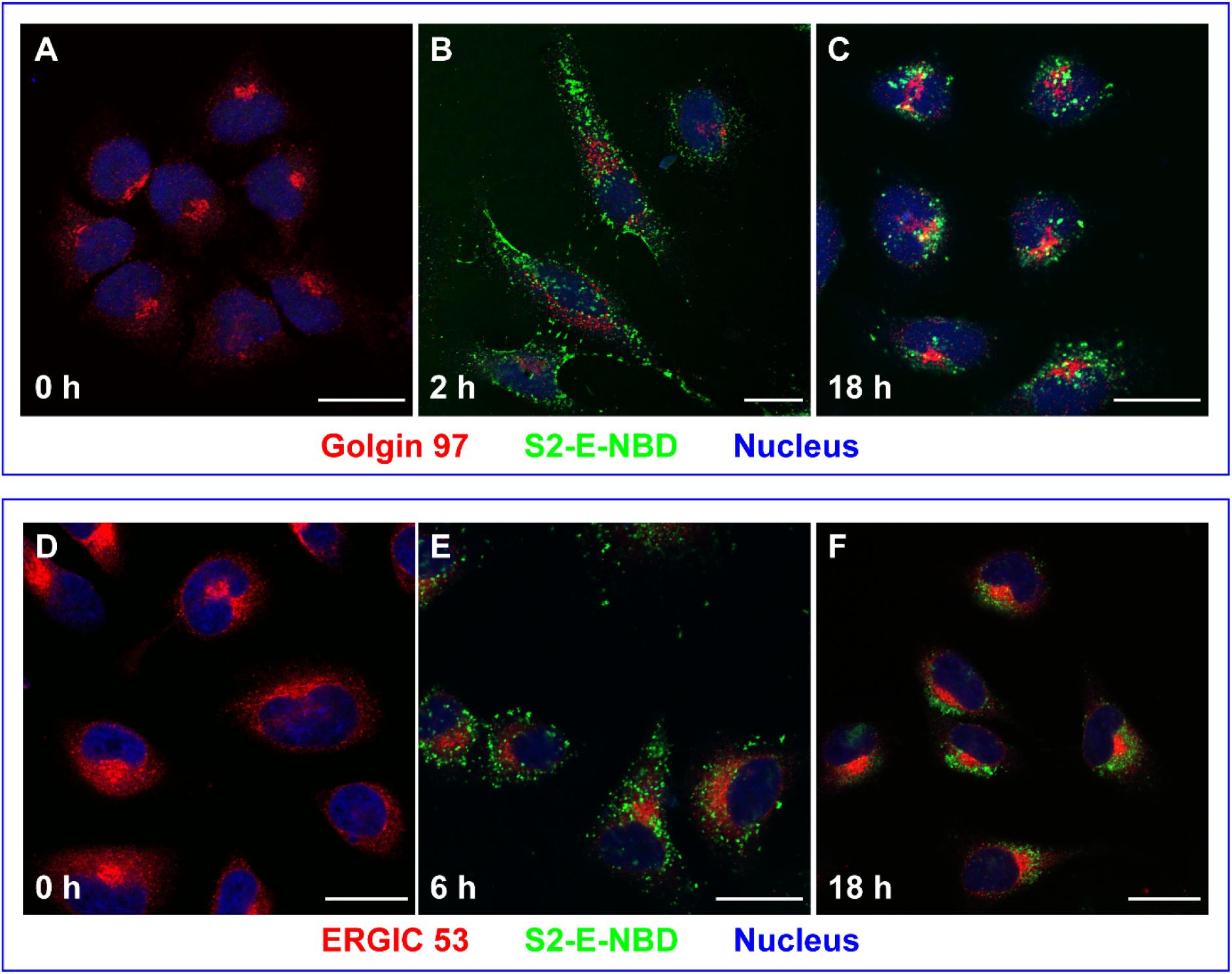
SARS-CoV-2 envelope protein traffics to a perinuclear location and accumulates near the Golgi and ERGIC compartments. Representative confocal microscopy images showing HeLa cells treated with 2.5 μM S2-E-NBD. Color markers are: green, S2-E labeled with NBD; red—in panels **A-C**—is from an antibody to Golgin-97, a Golgi marker; in panels **D-F**, red is from an antibody to ERGIC-53, a defining marker for the ERGIC region; blue is the fluorescent dye DRAQ5, marking the cell nucleus. Panels **(A)** and **(D)** are the control samples where cells were not treated with S2-E-NBD. Other panels are labeled with time following S2-E-NBD addition to the cell culture. Experiments were repeated 3 times using 3 different S2-E-NBD preparations. All scale bars are 25 μm. Further details in materials and methods and Supporting Information **Fig. S6**

It is likely that the Golgi-localization motifs (26) in S2-E drive its retrograde trafficking in a way closely related to the mechanism that facilitates E protein Golgi/ERGIC retention during viral infection. However, we cannot rule out the possibility that the retrograde trafficking documented in **Fig. 3** reflects the outcome of a cellular stress response to amphipol-delivered S2-E. Isolated coronavirus E overexpression in transiently transfected model mammalian cell lines is known to induce apoptosis (42,43). However, comparative studies of cell infection with SARS versus SARS lacking the E gene have shown that lower levels of E protein can modulate the unfolded protein response (UPR) and thereby mitigate apoptosis (44). It is plausible that the amphipol-mediated extracellular delivery of S2-E triggers cell stress and UPR-related retrograde trafficking, leading to deposition of S2-E in perinuclear aggresomes. Aggresomes are ordered protein aggregates that form following transport of certain proteins along microtubes by dynein to perinuclear microtubule-organizing centers (45). Interestingly, previous reports have linked aggresome formation and their subsequent clearance via autophagy to coronavirus replication (46–48). Further study is clearly required. For now, we can confidently state that delivered S2-E ultimately traffics back to a perinuclear area that is immediate to the Golgi and ERGIC compartments which mirrors the localization of SARS-CoV-2 infected cells.

## Conclusions

We have shown that the S2-E protein can be stripped of lipid and detergent and purified into aqueous solutions in which its solubility is maintained solely by complexation with amphipols. The protein can then be delivered to lipid bilayers, in which the protein spontaneously inserts into the membrane to form ion channels. Likewise, addition of the S2-E protein to living human cells results in plasma membrane integration and subsequent retrograde trafficking deep within the cell to a location immediately adjacent to both Golgi and ERGIC compartments, which are believed to be the key locales of coronavirus replication and assembly. The S2-E protein-to-cells approach established by this work should be exploitable as a route to delivering chemically modified full length S2-E to cells in culture or possibly even to cells under physiological conditions. This capability enables a wide range of chemical biological tools to explore the biological function of this protein or to test whether chemical warhead-armed S2-E can play the role of a Trojan horse to interfere with SARS-CoV-2 replication, potentially as an anti-COVID therapeutic or prophylactic. The results of this work also establish a general paradigm for using amphipols to deliver membrane proteins to living cells, although whether numerous other membrane proteins can be successfully delivered using this approach remains to be explored.

## Data Availability

All data needed to evaluate the conclusions in the paper and supporting information are presented in the manuscript or in the supporting information. Correspondence and requests for materials should be addressed to WDVH or CRS.

## Acknowledgements

Special thanks to Abigail C. Neininger for help with fluorescence microscopy.

## Author Contributions

JMH, RC, DDL, and AH conducted all experiments for this work. All authors participated in data analysis and wrote the paper. WDVH and CRS conceived of the work and directed the approaches used.

## Abbreviations

(SARS-CoV-2): Severe acute respiratory syndrome 2 virus
(S2-E): SARS-CoV-2 envelope protein
(ERGIC): endoplasmic reticulum-to-Golgi intermediate compartment
(CoV): coronavirus
(VLP): virus-like particle
(ARDS): acute respiratory distress syndrome
(NBD): nitrobenzoxadiazole
(UPR): unfolded protein response

