## Supporting Information for "Delivery of recombinant SARS-CoV-2 envelope protein into human cells"

**Contents**

**Supporting Materials and Methods**

**Supporting References**

**Supporting Figures S1-S6**

### Supporting Materials and Methods

#### Recombinant SARS-CoV-2 Envelope protein construct

DNA encoding the SARS CoV-2 envelope protein (S2-E) sequence followed by a C-terminal linker with a thrombin cleavage site and 10X His tag was inserted into a pET-21b plasmid vector. The encoded S2-E sequence is as shown below, with the added linker/thrombin site/tag indicated in blue font:

MYSFVSEETGTLIVNSVLLFLAFVVFLLVTLAILTALRLCAYCCNIVNVSLVKPSFYVYS-
RVKNLNSSRVPDLLVLESSGGGSILVPRGSGGSHHHHHHHHHH

Multiple constructs were tested for expression in several bacterial strains and the inclusion of the linker with thrombin cut site significantly increased overexpression levels.

#### Recombinant expression of the SARS-CoV-2 Envelope protein

The expression plasmid indicated above was transformed into Rosetta 2(DE3) pLysS cells. Transformed cells were then spread on agar LB plates supplemented with ampicillin and chloramphenicol before overnight growth at 37 °C. Colonies were picked the next afternoon and used to inoculate LB medium (150 mL in 500 mL baffled flask) supplemented with ampicillin and chloramphenicol before culturing the bacteria overnight with rotary shaking at 37 °C at 230 rotations per minute (RPM). The next morning, 20 mL of overnight culture was added to 1L of antibiotic-supplemented M9 medium with added M9 salts

(Corning) in a 2L baffled flask and grown at 37 °C at 230 RPM. After ~ 5-8 hours, when the cultures had grown to an $OD_{600}$ of 0.6-0.8 they were induced with 1 mM isopropyl β-D-1-thiogalactopyranoside. Induced cultures were allowed to incubate overnight at 37 °C at 230 RPM to facilitate production of S2-E inclusion bodies. Cultures were harvested the next morning by spinning cultures down at 3500 x g for 20 min at 4 °C and the pelleted cells were flash frozen in liquid nitrogen before being stored at – 80 °C. Six liters of culture typically resulted in a wet cell mass of 10-14 g.

**Purification of the SARS-CoV-2 Envelope protein**

10-14 grams of frozen cells were thawed at room temperature and suspended in 140 mL of lysis buffer {75mM tris(hydroxymethyl)aminomethane (Tris) pH 7.8, 300 mL NaCl, and 0.2 mM ethylenediaminetetraacetic acid (EDTA)}, supplemented with 0.5 mM magnesium acetate, lysozyme, ribonuclease, deoxyribonuclease, and 50 µL/gram wet cell mass protease inhibitor (P8849 Sigma). The lysis slurry was tumbled for two hours at 4 °C before being sonicated on ice for 12 minutes at 60 % power with alternating 5 second pulses and pauses (total power imparted on the slurry was ~ 80 kJ). The sonicated slurry was centrifuged at 25,000 x g for 20 minutes at 4 °C and the supernatant was discarded. The inclusion body pellets were resuspended with a Dounce homogenizer in 140 mL of lysis buffer. The sonication and centrifugation steps were repeated once in order to further clean the inclusion bodies. Clean inclusion body pellets were resuspended in 140 mL of lysis buffer supplemented with 3 % w/v N-dodecyl-N,N-dimethylglycine (Empigen), 0.5 mM dithiothreitol (DTT), and 25 µl/gram wet cell mass protease inhibitor before being tumbled overnight at 4 °C. The next morning, persistently insoluble materials were removed from the dissolved inclusion bodies by spinning at 25,000 × g for 45 minutes at 4 °C. The supernatant was saved.

2 mL of HisPur Ni-NTA Superflow agarose resin (Thermo Scientific) was equilibrated in lysis buffer before being mixed with the supernatant. The resin and supernatant were tumbled 1-2 hours at 4 °C before the resin was loaded onto a gravity column connected to an $A_{280}$ detector. To wash away unbound impurities, 15 column volumes (CV) of 0.3% Empigen, 1×Tris-buffered saline (TBS, 20 mM Tris pH 7.5 140 mM NaCl), plus 0.25 mM DTT were passed through the resin. Low-affinity impurities were washed away with 15 CV washes of 30 mM imidazole, 1× TBS, 0.1% 1-myristoyl-2-hydroxy-sn-glycero-3-phosphocholine (LMPC), and 0.25 mM DTT. This was repeated for 75, 90, and 120 mM imidazole washes in order to remove the remaining impurities. A 10 CV solution of 0.2 wt% PMAL-C8 (Anatrace, Maumee, OH) in 1X TBS was washed over the resin in 2 CV pulses in order to remove the majority of the detergent and exchange the resin-bound envelope protein into amphipols. Excess amphipols and residual detergent were then removed by washing the column with 10 CV of 1× TBS. Amphipol-complexed envelope protein was then eluted from the column with 5 CV of 250 mM imidazole in 1× TBS pH 7.8. It was seen that envelope protein could be eluted in either amphipol PMAL-C8 or A8-35 (Anatrace), but PMAL-C8 was chosen because it is zwitterionic and easier to work with than the anionic A8-35. The S2-E construct tag can be proteolytically removed with thrombin in detergent solutions for structural and biochemical studies, this was not carried out here because the thrombin precipitated the S2-E in PMAL-C8 amphipols solutions.

**Fluorescent labeling of S2-E**

Immediately after the envelope protein was eluted in complex with PMAL-C8, 1.5 mg of thiol-reactive N,N'-dimethyl-N-(iodoacetyl)-N'-(7-nitrobenz-2-oxa-1,3-diazol-4-yl)ethylenediamine (IA-NBD) (Thermo Fisher) was dissolved into 300 µL of dimethyl sulfoxide (DMSO) and added to the ~10 mL of eluted protein for labeling of one or more of its 3 cysteine residues (which are sequentially proximal within the S2-E extramembrane C-terminus). The reaction tube was covered in foil and tumbled at room temperature for one hour before being filtered through a 0.8 µm Acrodisc low-protein binding filter (PALL PN 4618) . The filtered amphipol/S2-E solution was then extensively dialyzed against 1× TBS pH 7.8 with 0.25 mM tris(2-carboxyethyl)phosphine hydrochloride (TCEP) and EDTA in 6-8 kD molecular weight cut-off (MWCO) dialysis tubing from Spectra/Por (part number 132660). On the morning of tissue culture treatment, the dialyzed protein was concentrated ~ 10 X in a 10kD MWCO Amicon Ultra-15 filter cartridge by centrifuging at 2500 x g at room temperature (RT). Protein purity was checked by sodium dodecylsulfate polyacrylamide gel electrophoresis (SDS-PAGE) and the concentration was determined by measuring absorbance at 280 nm using an extinction coefficient of 6000 $M^{-1}cm^{-1}$. Dye labeling efficiency was checked by measuring absorbance at 472 nm with an excitation coefficient of 23700 $M^{-1}cm^{-1}$. It was seen that the labeling efficiency was usually ~0.7 NBD per protein, indicating that, on the average, only one of three wild type cysteine residues was modified. NBD-modified S2-E (S2-E-NBD) was normally used within one week (with storage at 4 °C) following completion of preparation. Indeed, we observed that preparations that were stored for longer than 3 weeks at 4 °C led to cellular results in which S2-E-NBD was seen to traffic as usual to the perinuclear space but did not then obviously segregate to one side of the nucleus at longer timepoints (16/24h), as was seen when freshly prepared samples were employed. Starting with six liters of culture, this protocol yields roughly 500 µg of fluorescently labeled S2-E protein.

**Functional delivery of SARS-CoV-2 envelope protein from amphipols to planar bilayers**

Planar lipid bilayers were formed from a solution of synthetic 1-palmitoyl-2-oleoyl-glycero-3-phosphocoline (POPC) and 1-palmitoyl-2-oleoyl-glycero-3-phosphoethanolamine (POPE, Avanti Polar Lipids) in a 3:1 mole ratio in *n*-decane (Sigma-Aldrich). A glass bulb was used to apply the POPC/POPE solution in a 150 µM aperture of a Delrin cup (Warner Instruments), separating the *cis* and *trans* chambers. Asymmetric buffers were used, with 5 mM 4-(2-hydroxyethyl)-1-piperazineethanesulfonic acid (HEPES), 500 mM NaCl, pH 7.2 for the *cis* chamber, and 5 mM HEPES, 50 mM NaCl, pH 7.2 for the *trans* chamber as done in previous studies for comparison.(1,2) The bilayer capacitance ranged between 50-65 pF.

Delivery of the SARS-CoV-2 envelope protein (S2-E) complexed by PMAL-C8 amphipols to pre-formed planar bilayers was achieved spontaneously by pipette addition of ~1 µL of ~3 µM of the S2-E–amphipol complex and accompanied by stirring of the chambers. Each chamber (cis/trans) had 2.5 mL of buffer (5mL total volume). Incorporation of S2-E was achieved by addition of PMAL-C8 complexed S2-E to either the cis or to both cis and trans chambers. The final concentration of protein varied from ~5 nM–15 nM. S2-E unitary currents were collected from a BC-535 amplifier (Warner Instruments) at 23 ± 1°C, 10 mV/pA gain and low-pass filtered by a 4-pole Bessel filter at 1 kHz. The resulting data were digitized using an analog-to-digital converter (Digidata 1550A, Molecular Devices) at 1 kHz using the pClamp10.7 software suite (Molecular Devices). The single-channel data were collected over separate amphipol delivery experiments carried out on different days, with three-minute recordings for each voltage. Recordings were analyzed using the single-channel search in ClampFit 10.7 (Molecular Devices) filtered with a 100 Hz low pass Gaussian filter and ignoring short level changes of 25 ms duration to obtain the number of events, open probability, reversal potential, and single-channel conductance.

**Maintenance of cell cultures**

Human cell lines HeLa (ATCC cat# CCL-2) and alveolar SW1573 (ATCC cat # CRL-2170) cells were obtained from ATCC. HeLa cells were maintained in a tissue culture incubator with humidified air supplemented with 5% $CO_2$ at 37°C. HeLa cells were grown in low glucose Dulbecco's Modified Eagles Medium (DMEM, Gibco cat#11885-084) supplemented with 10% fetal bovine serum (FBS, Gibco 26140-079), 1% penicillin/streptomycin (P/S). SW1573 cells were kept in a tissue culture incubator with humidified air without added $CO_2$ at 37°C. The SW1573 were grown and maintained in (Leibovitz's medium # 15, or L-15; ATCC cat # 30-2008) supplemented with 10% FBS and 1% P/S. Both cell lines were passaged 2 times a week and seeded such as to remain below 100% confluency.

**Cell culture treatments**

The underlying idea our S2-E-to-cell experiments was to treat cells with aliquots of fluorescently-tagged S2-E/amphipol and then to follow cell association and intracellular trafficking of the flurorescently-taged protein over time. These experiments were conducted with healthy growing cells that are well attached to a glass coverslip surface. These cells were treated with various concentrations of SE-2 protein/amphipol for differing amounts of time. This is achieved with freshly concentrated, and preferably freshly made, SE-2 protein complexed with amphipol PMAL-C8 in TBS. In order not to perturb the cells by changes in osmotic pressure, pH, or temperature the stock S2-E amphipol solution was processed prior to addition to the cells. S2-E was first dialyzed against TBS. Further, the resulting S2-E/amphipol solution was added to fresh 37°C pre-warmed cell media and mixed before aliquots were then added to cells. The added volume of the S2-E/amphipol solution added to the cell culture never exceeded 10% of the culture volume and was often less (4-6%). At the chosen time points, the coverslip containing attached cells was rinsed to discard floating cell debris and then 'fixed' with paraformaldehyde. Fixing is intended to chemically freeze cells. The procedure kills the cells but preserves their plasma membrane surface and intracellular structures. After discarding excess fixative, cells are treated differently depending on the cell structures we wished to observe in relation to S2-E protein. When the goal was to see S2-E in relation to the plasma membrane we used lecithin conjugated to a fluorophore together with a cell nucleus dye DRAQ5 which preferentially binds double-stranded DNA. When the intent was to observe intracellular organelles Golgi and ERGIC, the fixed cells were permeabilized by incubation with detergent with the goal being to make small holes of sufficient size for antibodies, specific to those structures, to enter the cell surface and gain access to Golgi and ERGIC.

Exponentially growing cells were plated in the wells of a 12-well plate with 1.5 thickness glass coverslips (Fisherbrand cat # 12-545-81) at a cell density of 25,000 live cells/well in 2mL cell media. After a day, the cells destined for 24h and 16-18h time points were treated by rinsing once with Dulbecco's phosphate buffered saline without calcium and magnesium (DPBS, Corning cat 21-031-CV) and by adding 1mL of fresh medium with no more than 5-10 % volume of freshly concentrated S2-E protein (NBD-labelled or not) with amphipol in the dialysis buffer at the concentrations stated in the figure captions. Cells were protected from light. The remaining and undiluted S2-E/amphipol stock was stored at 4 °C overnight protected from light and used in the same way the next morning for the remaining time points. At the appropriate time, each coverslip was transferred into a fresh 12-well plate well facing upwards containing 2 mL of 37 °C pre-warmed DPBS. DPBS was discarded by suction, and cell fixation was started by adding gently 1 mL/well of pre-warm 4% paraformaldehyde (PFA, EMS cat #15714) dissolved in DPBS for 15 min at 37°C. After fixation in 4% PFA, all steps were carried out with gentle mixing and at room temperature and with plates protected from light by being kept wrap in aluminum foil. All solutions were filtered prior to addition to coverslips to avoid addition or formation of aggregates. Fixation was stopped by discarding the 4% PFA solution by suction and by gently rinsing 2 times with 2mL/well with DPBS for 15 min/each. The cell membrane was labeled with wheat germ agglutinin conjugated to Alexa Fluor-555 or WGA-AF555 (Invitrogen cat# W32464) at a final concentration of 5µg/mL for 15min using 1mL/well. Coverslips were then rinsed from excess WGA-AF555 by rinsing 2 times at 2mL/well in DPBS for 15 min for each time point and once using 100mM glycine in DPBS. The times of cell treatment were imperfectly staggered. After the 100mM glycine rinse, the timepoint samples were set aside without shaking until other time points were completed. When all time points had finished the 100mM glycine rinse, all coverslips were transferred into a common 12-well plate, and the following treatments proceeded as a group.

For time-course S2-E-NBD trafficking course experiments without antibodies (Fig. 2 and S3) cells were cell nucleus-labeled using freshly opened of DRAQ5 (Invitrogen cat #65-0880), an anthraquinone membrane permeable dye with high affinity for double-stranded DNA. DRAQ5 was used diluted 1:2500 in DPBS using 2mL/well and incubated for 1h. Excess DRAQ5 was then rinsed away twice with 2mL/well 10min each DPBS +0.01% Triton X-100 (TX-100) and once with DPBS only. Coverslips were mounted in glass slides using Prolong Gold antifade (Invitrogen, cat # P10144) and cured, typically for 2 days, before imaging.

For experiments requiring antibody detection of organelle-specific marker proteins (Fig. 3 and S3), cells were permeabilized with 1mL/well containing 0.1% TX-100 dissolved in DPBS for 15min. Permeabilization was stopped by rinsing twice in 2 mL/well DPBS and blocking overnight with 2mL/well of 1% BSA dissolved in DPBS with 0.01% TX-100. The day after, plates were allowed to reach RT for 30-40 min without mixing and then treated with specific antibodies for detection of Golgin97 (1mL/well, diluted 1: 500; Invitrogen, cat #A21270), or ERGIC53 (3) (1mL/well, diluted 1:500; ENZO cat# ENZ-ABS300), or for E protein detection (1:1000 dilution of rabbit polyclonal antibody ProSci cat# 10-518) diluted in freshly made and 0.2µm filtered 1% BSA dissolved in DPBS + 0.01% TX100 (1%BSA) for 2h at RT with gentle mixing. Each coverslip was treated to remove unbound primary antibody by rinsing 3 times with DPBS + 0.01% TX-100, 15 min each. Samples were then incubated with a secondary antibody conjugated to a fluorescent dye using 1mL/well for 1 hr in 1% bovine serum albumin (BSA) using 1:1000 dilution of goat anti-mouse conjugated to Alexa Fluor 546 (AF546; Invitrogen cat# A11030), mouse mAb for Golgin-97 and ERGIC53, or donkey anti-rabbit conjugated Alexa Fluor 488 (AF488; Invitrogen cat# A32790) for samples treated with rabbit anti-E polyclonal. After 1 h samples were rinsed twice for 15min with 2mL/well DPBS + 0.01% TX100 and treated in the same way as above for nuclear counter-staining using DRAQ5 and mounting.

**Cell imaging**

All experiments shown were imaged using a LSM 510 confocal microscope, with a Plan-Neofluar 40x/1.30 Oil DIC objective. The confocal pinholes were set at 84µm for the 633 nm HeNe1 laser line, at 80µm, for the 543 nm HeNe1 laser line, and at 86µm for the Argon laser. These settings correspond to a pinhole diameter of 1.00 Airy units for the HeNe2 laser (633 nm), and the same optical slice of 1.1µm for all three lines. These settings were kept for all channels in all experiments. The fluorophores were excited using the 488 nm line of a 40 mW Argon laser set at 10% power (reduced at source to 50%, for the final 5%) for NBD and AF488 signal. The 543 nm line was set at 10% power for WGA-AF555 and for anti-mouse AF546, whereas the 633 nm line of a HeNe laser was set to 15-50% power for DRAQ5 depending on the dye emission intensity. Images were collected at 1-2x and occasionally 3x zoom. The frame size was set usually to 1024, data depth was 8 bit, and Multi track/frame mode was used, for each channel. The scan average parameter was usually set to 4, but sometimes also 8 or 16. For presentation purposes, images were processed using ImageJ software.

**Statistics**

The numbers of replicates, validation of reagents, software, and statistical approaches are detailed in the above methods and/or in the relevant figure captions.

**Supporting Figures**

**Figure S1**

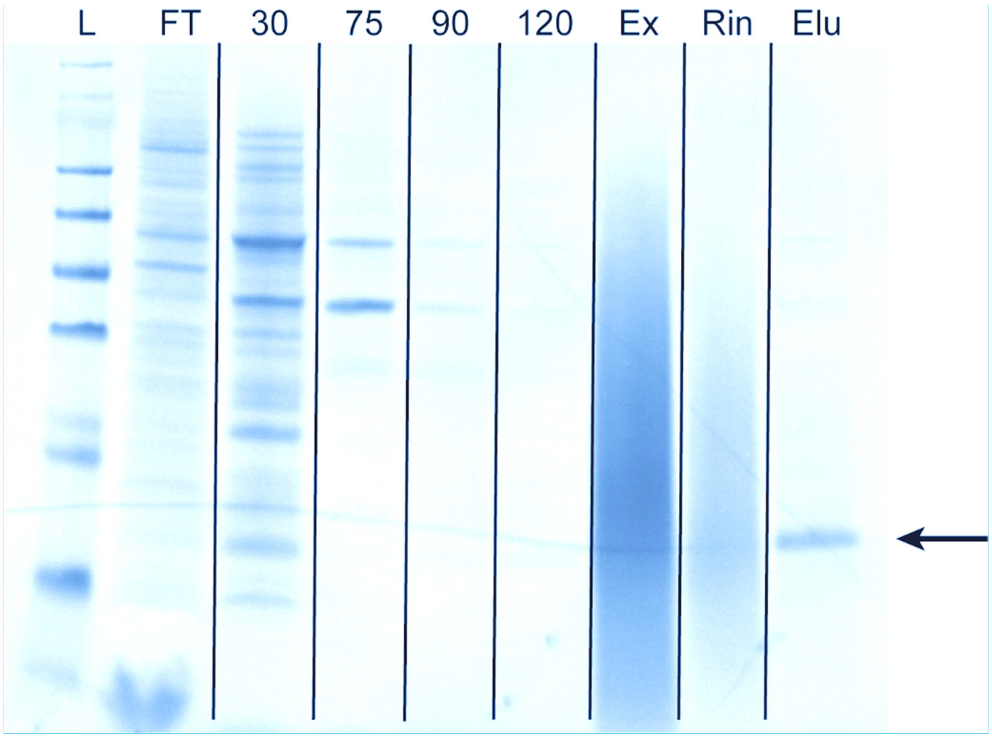


**Figure S1. Representative SDS-PAGE of SARS-CoV-2 purified into PMAL-C8.** Each lane represents a step in the purification protocol after passing detergent-solubilized inclusion bodies over the column. SDS-PAGE samples were boiled and reduced with 5mM DTT before being run. Gels were stained with the SimplyBlue coomassie alternative (Invitrogen). Lanes from left to right are: Molecular Weight Standard **L**adder SeeBlue Plus2, **F**low-**T**hrough, **30** mM imidazole wash, **75** mM imidazole wash, **90** mM imidazole wash, **120** mM imidazole wash, **Ex**change into PMAL-C8, **Rin**se unbound PMAL-C8, and **Elu**tion. The black arrow points to the purified S2-E protein in the final elution.

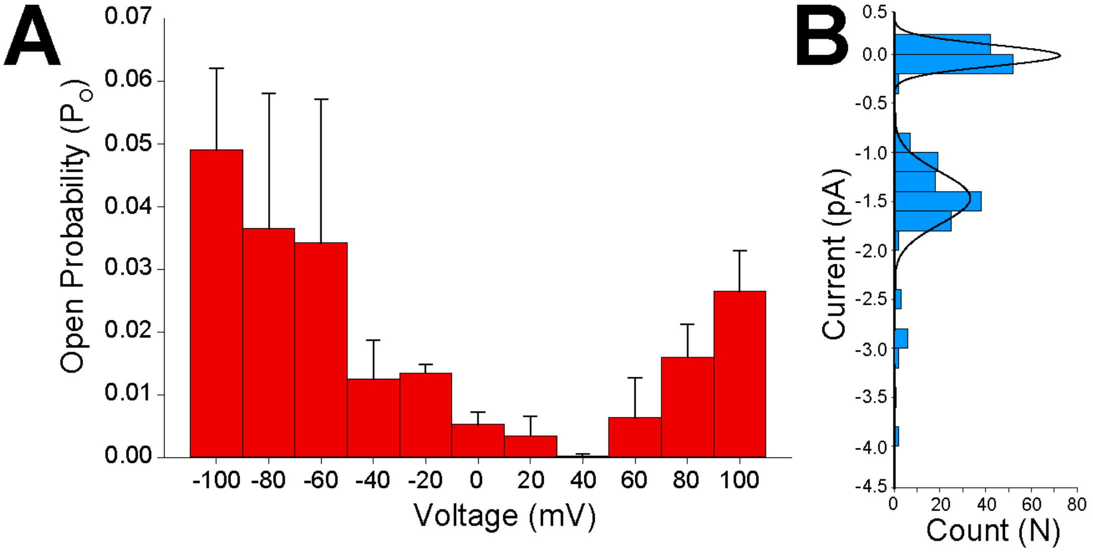


**Figure S2. S2-E ion channel open probability and current amplitude distribution from planar bilayer electrophysiology measurements after amphipol delivery.** Previous studies of CoV E proteins indicate that the channel has low open probabilities and current amplitudes. **(A)** Shows the open probability as a function of voltage for the data in Fig. 1*C*. Error bars are SEM from three distinct delivery experiments on different days and in total correspond to 9 min of measurement. **(B)** Identifies the number of events and the respective currents recorded at -100 mV. The historgram was fit to a 3 polynomial Gaussian fit with an average amplitude of 1.47 ± 0.05 pA for the open state. As in panel A, these data are from three separate recordings.

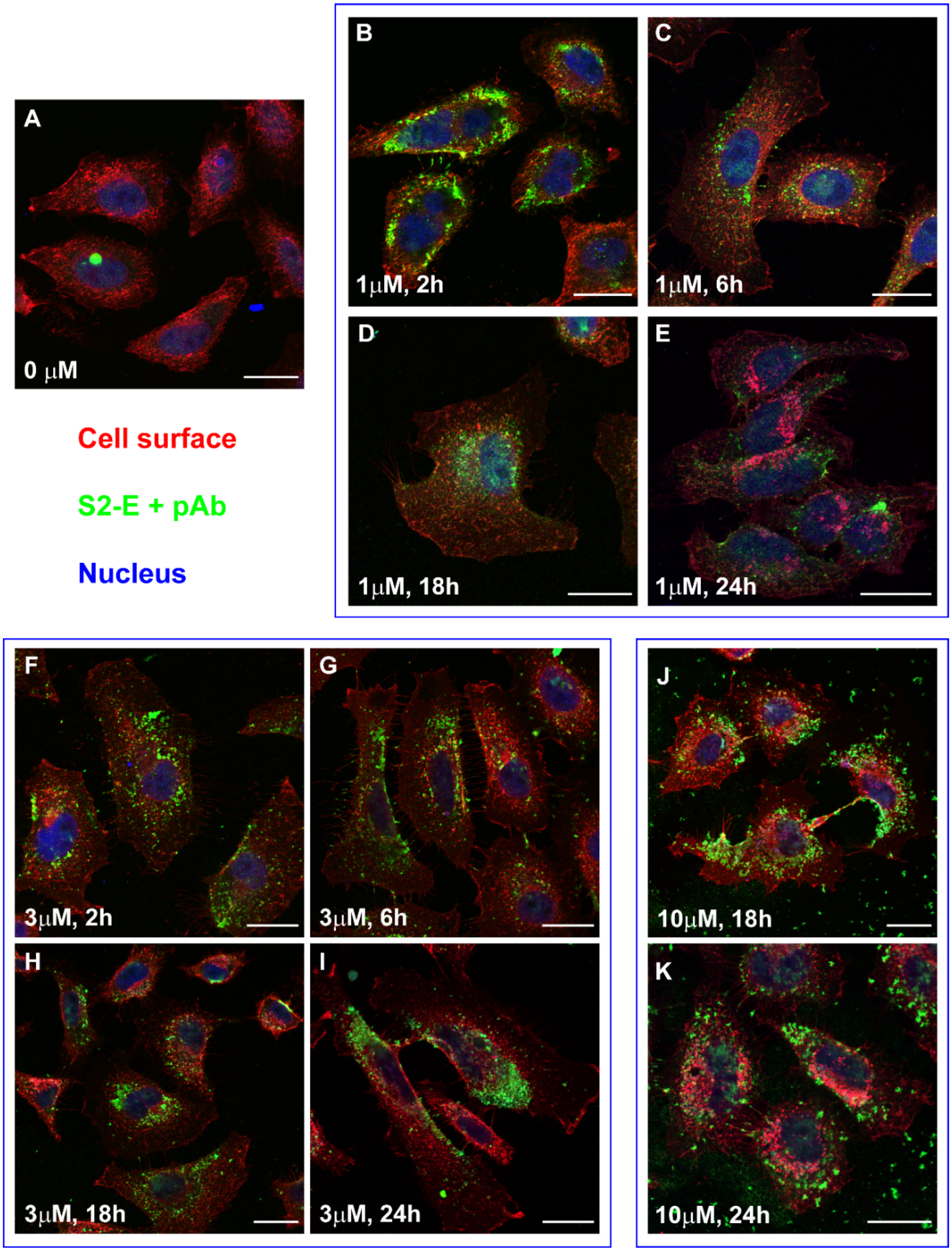
A
0 μM
Cell surface
S2-E + pAb
Nucleus
B
1μM, 2h
C
1μM, 6h
D
1μM, 18h
E
1μM, 24h
F
3μM, 2h
G
3μM, 6h
H
3μM, 18h
I
3μM, 24h
J
10μM, 18h
K
10μM, 24h

Figure S3. Lower Panel I. Individual channels and composite of upper panels A and B-E

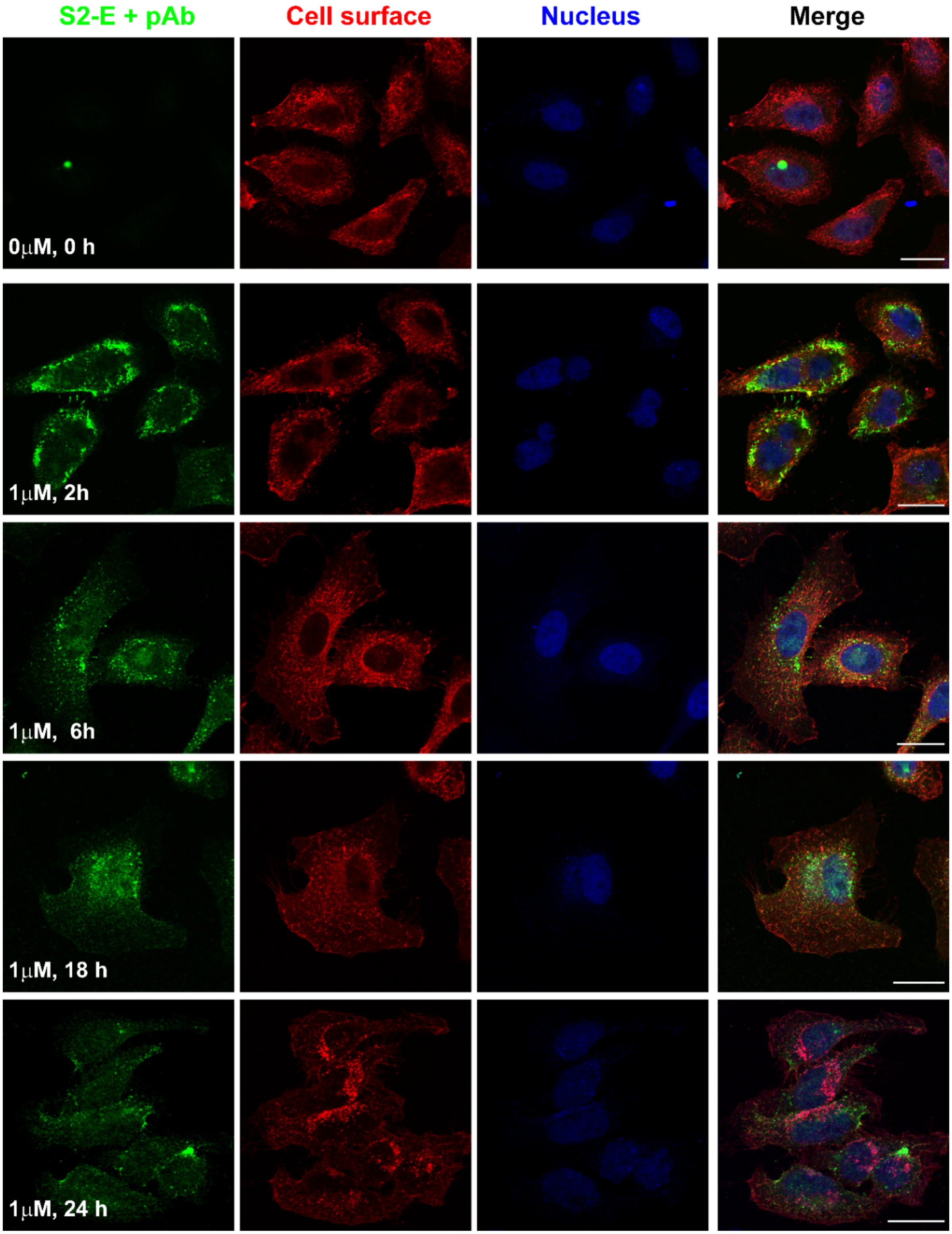

Figure S3. Lower Panel II. Individual channels and composite of upper panels A, F-I

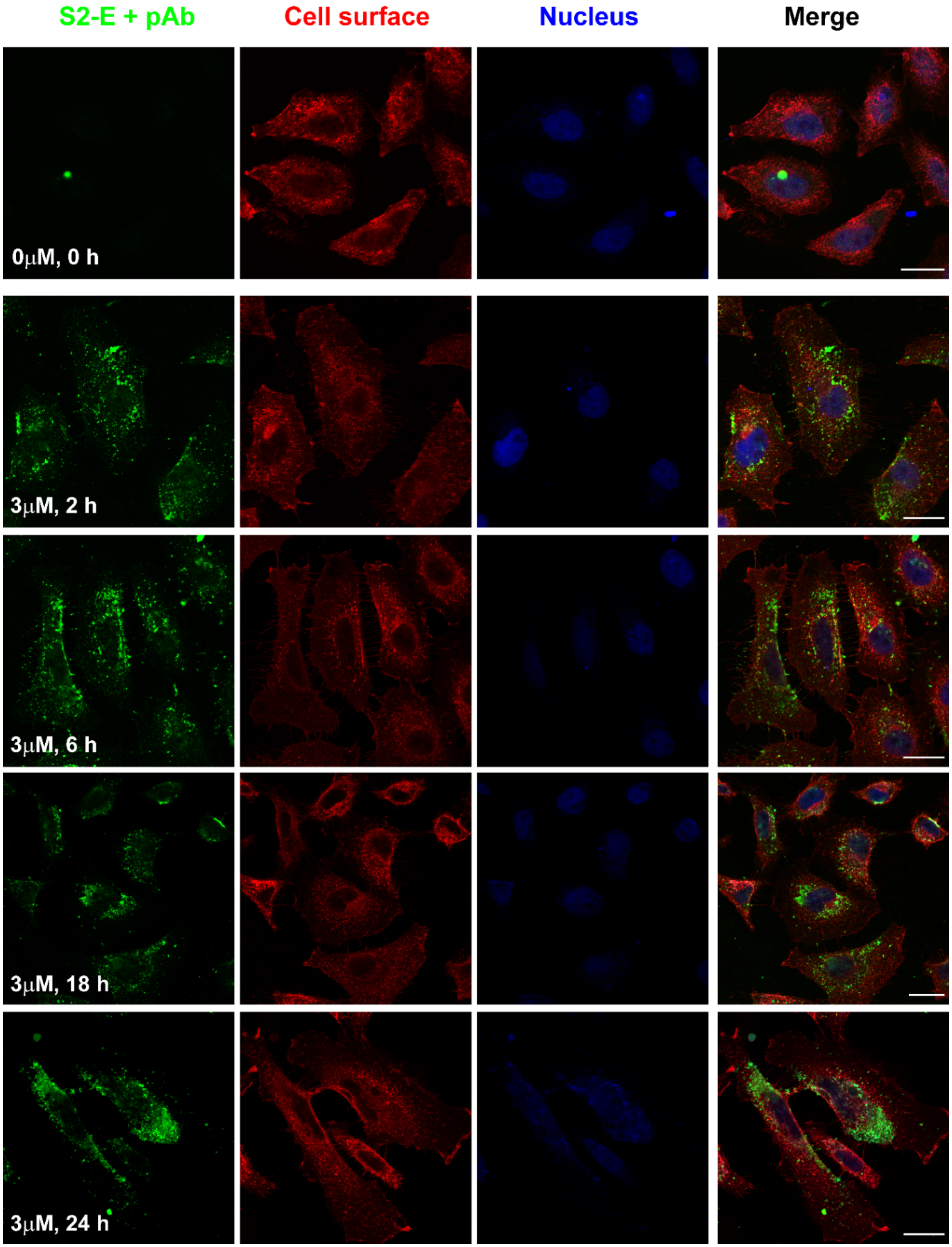

**Figure S3. Lower Panel III. Individual channels and composite of upper panels A and J-K**

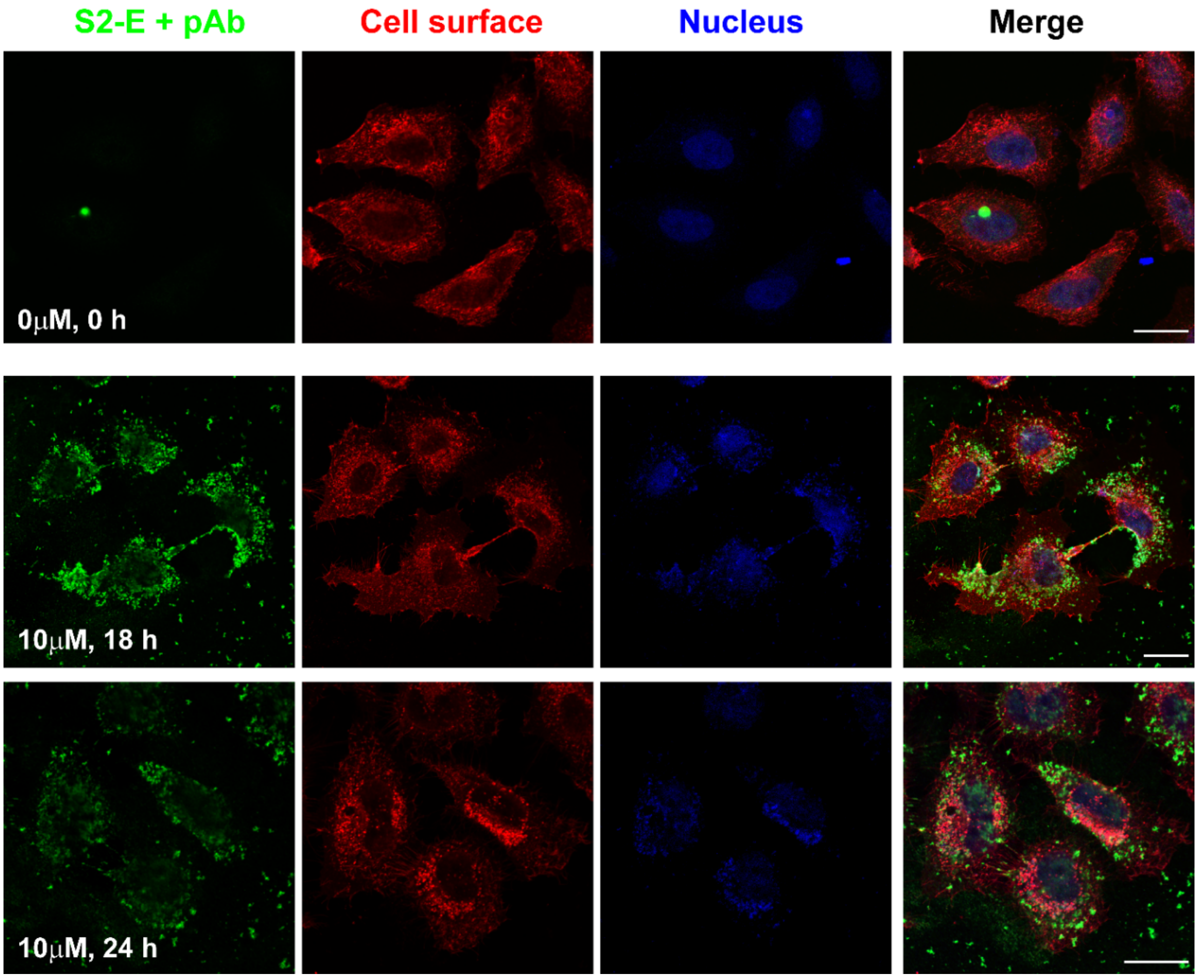


**Figure S3 (This page and 3 preceding pages). Unmodified S2-E protein delivered to cells from PMAL-C8 amphipol shows similar cell membrane localization and retrograde kinetics as for the NBD-tagged S2-E protein.** The time course and concentration dependency of unmodified S2-E was followed using rabbit anti-E polyclonal antibodies. Rabbit antibodies were detected using anti-rabbit conjugated AF488 (green). Hela cell surface plasma membranes (red) were detected using WGA-AF555 while cell nuclei (blue) were labeled using DRAQ5. Applied unlabeled S2-E concentrations were 1 μM for panels B-E, 3μM for panels F-I, and 10μM in panels J and K. Time points are: 0h and untreated sample (A); for the 1μM series: 2h (B), 6h (C) 18h (D) and 24h (E); for 3μM series: 2h (F), 6h (G) 18h (H) and 24h (I); for cells treated with 10μM unlabeled E: 18h (D) and 24h (E). At 10μM S2-E observation of cell debris suggest cellular toxicity. Overall, delivery and retrograde transport of unmodified S2-E follows a similar time course as for NBD-labeled S2-E; however, the control sample without added E (panel A) did exhibit a weak unspecific binding on the green channel. The lower panels I ($2^{nd}$ page of Fig.) , II ($3^{rd}$ page), and III ($4^{th}$ page) of the figure show the individual channels of each composite image presented in the upper panel. For lower panel I ( A-D), lower panel II (A, G-I) and panel III (A, J and K). All scale bars are 25 μm. This experiment was independently repeated twice. For further experimental details see the above Supporting Materials and Methods

**Figure S4. (2 Page Figure)**

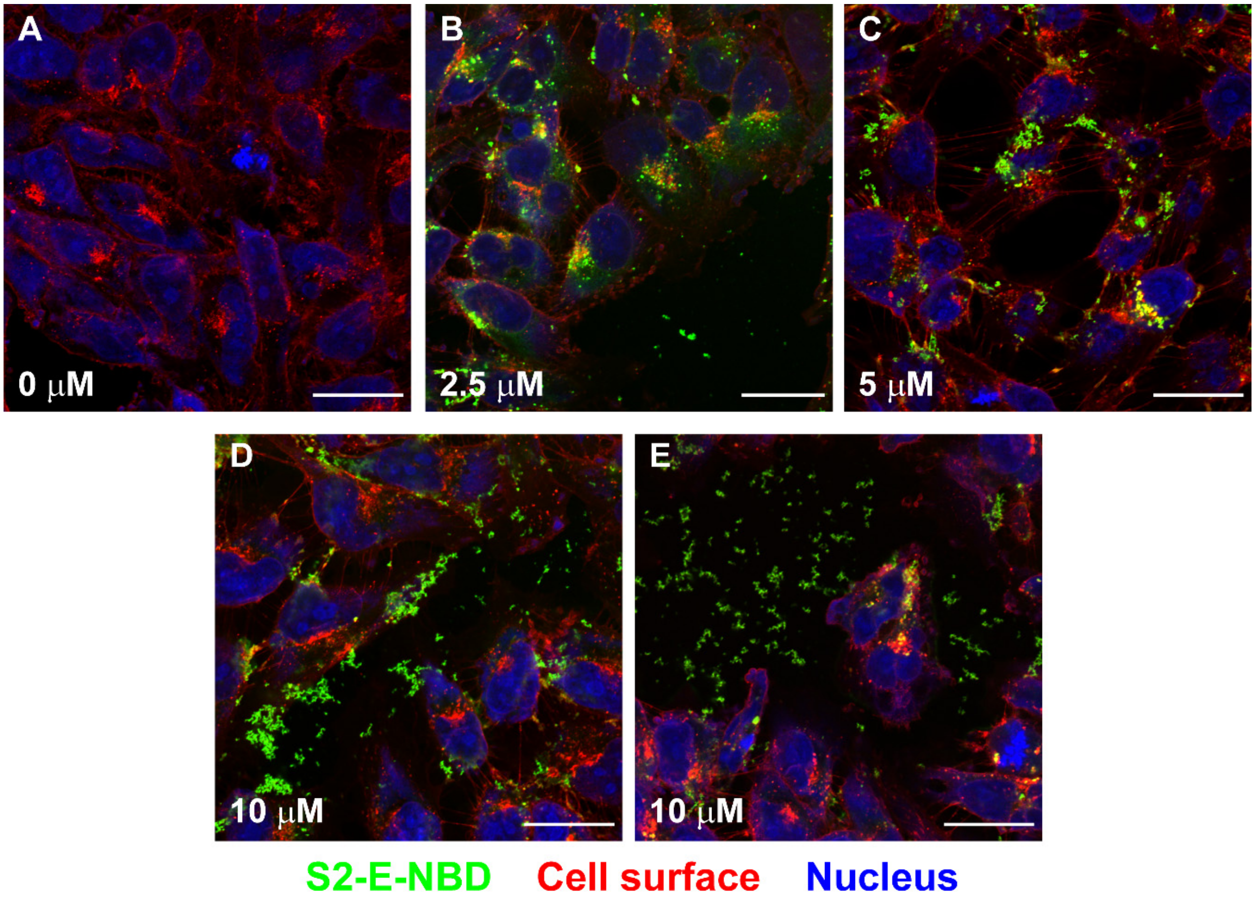

**Figure S4. Lower Panel. Individual channels and composite of upper panels A-D.**

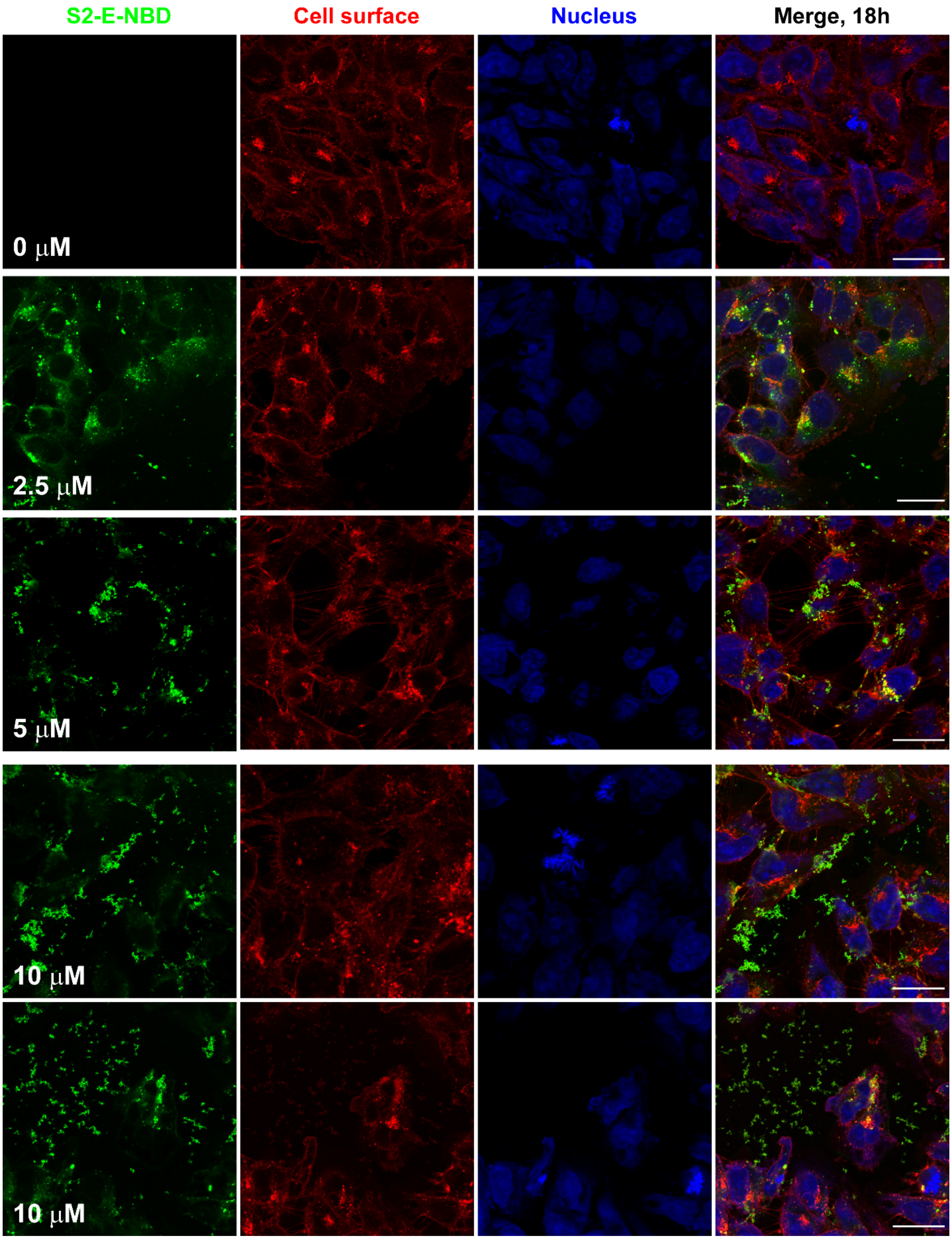

**Figure S4 (above 2 pages).** Cell membrane delivery and uptake of the SARS-CoV-2 envelope protein with PMAL-C8 amphipol in a human alveolar cell line. Representative confocal microscopy images showing SW1573 cells treated with increasing concentrations of added NBD-labeled S2-E. All time points were collected at 18h. **(A)** untreated control 0µM, **(B)** 2.5µM S2-E in the cell culture, **(C)** 5µM, and 10µM **(D and E)**. Color scheme is: green, NBD-labeled S2-E; red, cell plasma membrane (WGA-AF555); blue, cell nuclei (DRAQ5). As in HeLa cells, at 18hrs after cell treatment the S2-E-NBD signal is primarily at a perinuclear location (panels B and C). For cells incubated with the highest level (10µM) of S2-E-NBD, some the protein appears to be aggregated (D) while in panel E—top left side, SW1573 shows signs of co-localization of S2-E with the plasma membrane label, indicating cell membrane fragments, possibly suggesting cellular toxicity at 10µM S2-E-NBD. The lower panel (2$^{nd}$ page) of the figure shows the individual channels of each composite image presented in the upper panel. All scale bars are 25 µm. This experiment was conducted once. For further experimental details see the Supporting Materials and Methods.

Figure S5

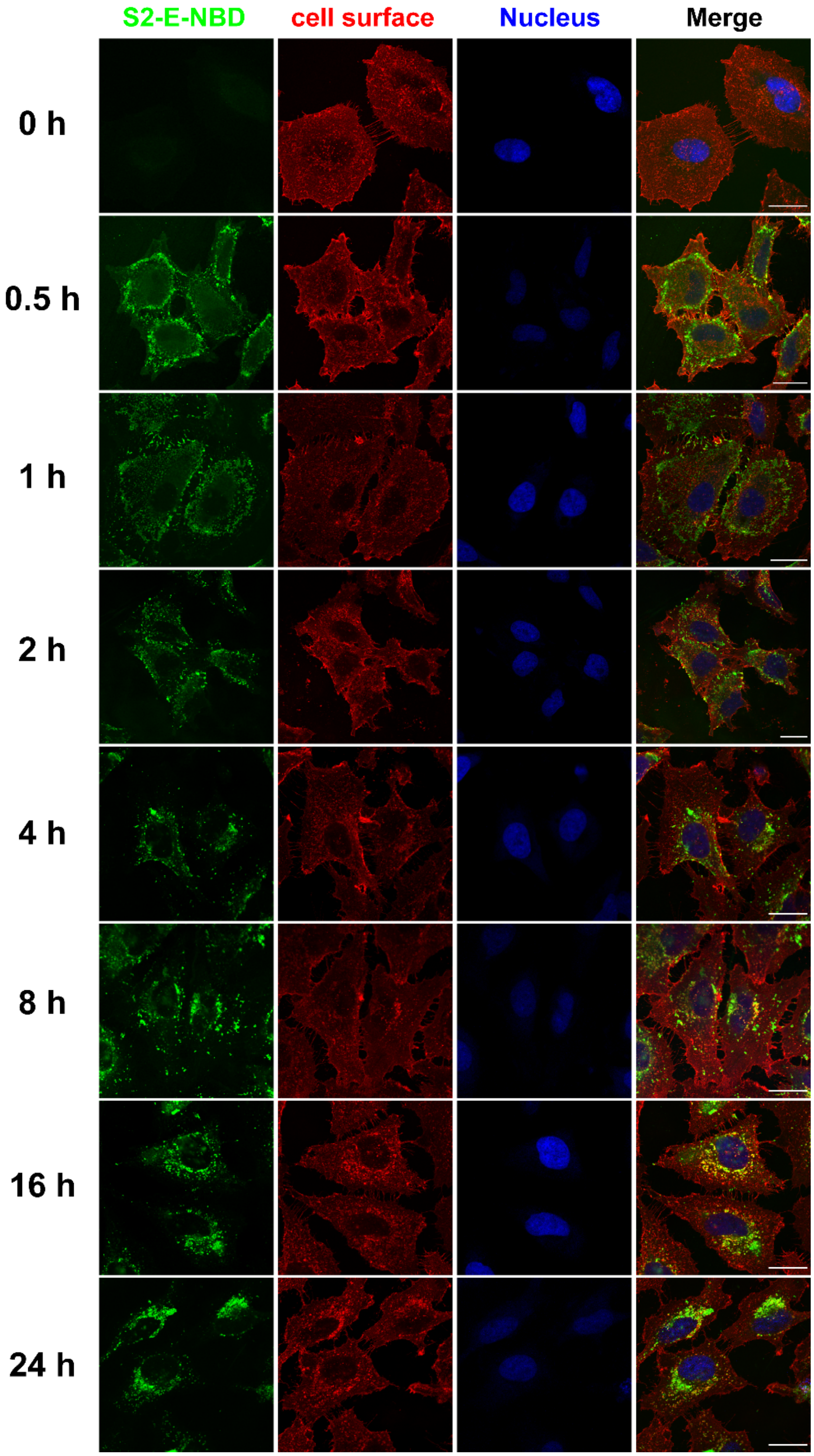

**Figure S5.** Individual channels for each composite image shown in Fig. 2 of the main paper. Color scheme: green, NBD-labeled S2-E; red, cell surface plasma membrane (WGA-AF555); and blue, cell nuclei (DRAQ5). Note the movement over time of the S2-E-NBD signal towards one side of the cell nucleus while the red signal remains constant.

Figure S6

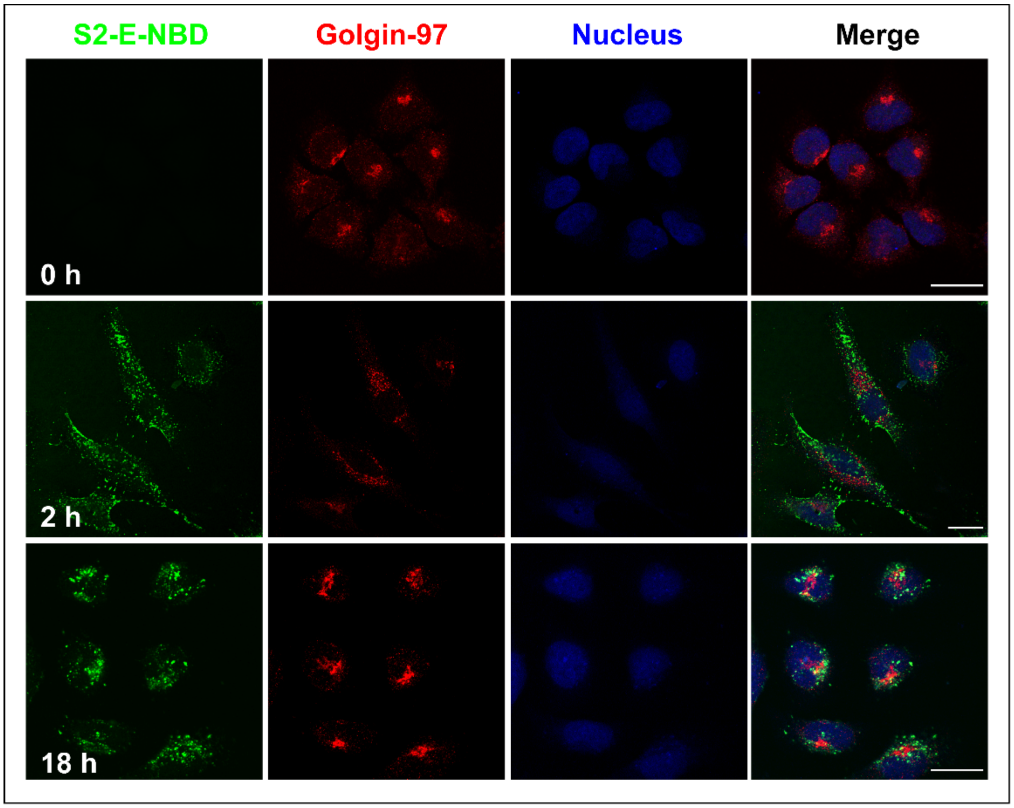
S2-E-NBD
Golgin-97
Nucleus
Merge
0 h
2 h
18 h

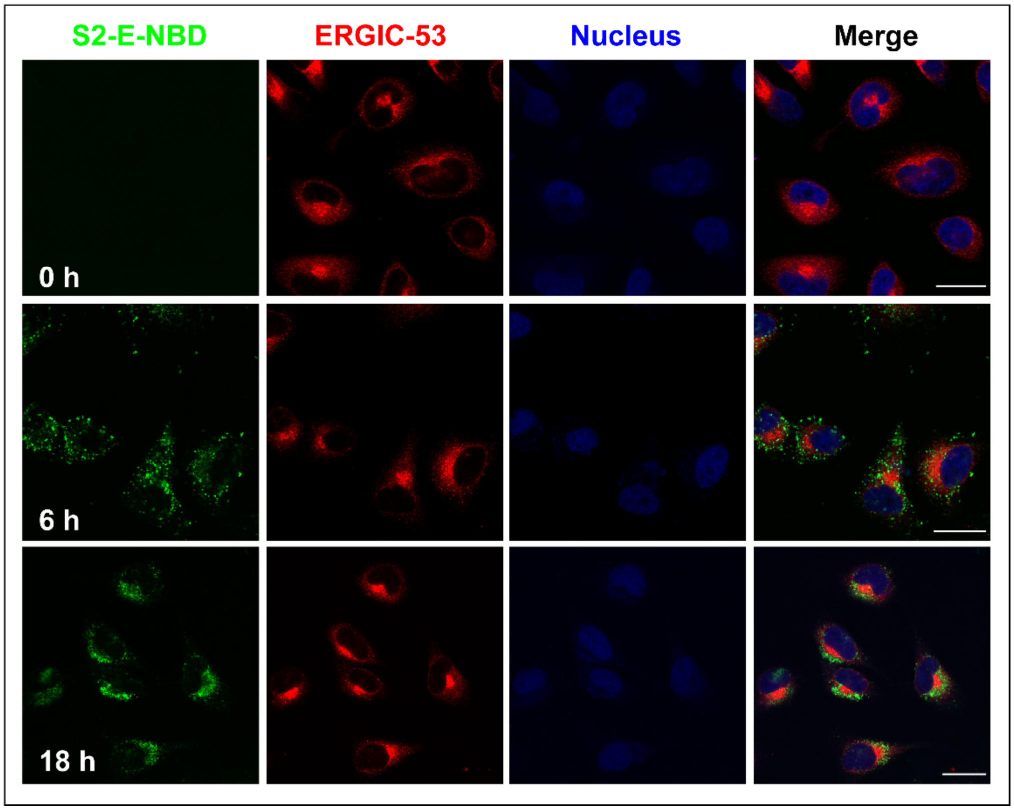
S2-E-NBD
ERGIC-53
Nucleus
Merge
0 h
6 h
18 h

**Figure S6.** The individual channels of each composite image shown in Fig. 3. Color scheme: green, S2-E label NBD and blue is cell nuclei in both panels, in the upper panel red is mouse monoclonal anti Golgin-97 a Golgi marker, while in the lower panel, red is the signal from mouse monoclonal antibody against ERGIC-53 an ERGIC-resident protein. Notice the movement of the S2-E-NBD signal over time toward the red signals from the Golgi marker. Also notice that some degree of overlap between red and green signals, indicating some degree of localization of S2-E at the Golgi.
